# All boreal forest successional stages needed to maintain the full suite of soil biodiversity, community composition, and function following wildfire

**DOI:** 10.1101/2022.11.18.517085

**Authors:** Teresita M. Porter, Emily Smenderovac, Dave Morris, Lisa Venier

## Abstract

Wildfire is a natural disturbance in boreal forest systems that has been predicted to increase in frequency, intensity, and extent due to climate change. Most studies tend to assess the recovery of one component of the community at a time but here we use DNA metabarcoding to simultaneously monitor soil bacteria, fungi, and arthropods along an 85-year chronosequence following wildfire in jack pine-dominated ecosites. We describe soil successional and community assembly processes to better inform sustainable forest management practices. Soil taxa showed different recovery trajectories following wildfire. Bacteria shared a large core community across stand development stages (~ 95-97% of their unique sequences) and appeared to recover relatively quickly by crown closure. By comparison fungi and arthropods shared smaller core communities (64-77% and 68-69%, respectively) and each stage appeared to support unique biodiversity. We show the importance of maintaining a mosaic ecosystem that represents each stand development stage to maintain the full suite of biodiversity in soils following wildfire, especially for fungi and arthropods. These results will provide a useful baseline for comparison when assessing the effects of human disturbance such as harvest or for assessing the effects of more frequent wildfire events due to climate change.

## Introduction

Climate change is one of the driving forces behind the anticipated increase in the frequency, intensity, and scale of wildfires in boreal regions.^1^ Wildfire is a natural stand-replacing disturbance in boreal forest systems affecting forest dynamics, biodiversity, and terrestrial carbon stocks that can reset ecological soil processes.^2^ Boreal forests are the second largest forest system after tropical seasonal forests and represent about 1.9 billion hectares in the northern hemisphere that are currently considered a net carbon sink.^3–5^ In Canada, jack pine (*Pinus banksiana*) is a common boreal forest species, source of timber, lumber, and pulpwood, and is prone to fire disturbance.^6^ As a result there are many fire-origin jack pine stands at various stages of establishment located in Canada. Understanding succession following natural disturbance regimes, especially for pyrophilous taxa, and the consequences of their perturbation due to climate change, is needed to mitigate these changes and adjust current forest management practices.^7,8^

The effects of wildfire on forested ecosystems are complex, affecting both the above ground and below ground components.^9^ Until now, most studies addressing boreal forest succession after wildfire have each focused on one of a relatively narrow suite of abiotic indicators of recovery such as soil nutrients, pH, fluxes in volatile organic compounds, or biotic indicators of recovery such as plants, arthropods, mosses, birds, or fungi.^2,4,10–16,17–20^ Environmental DNA sampling, coupled with high throughput sequencing and automated taxonomic assignment methods, also known as DNA metabarcoding, makes it possible to conduct large-scale biological surveys for biomonitoring.^21,22^ This approach, sometimes referred to as ‘biomonitoring 2.0’, has gained traction in government programs to improve scalability, reproducibility, and throughput, reducing the turnaround time between data gathering and reporting.^23–28^ Starting with a bulk environmental sample (e.g. homogenized soil), whole community DNA can be extracted without having to isolate or identify individual organisms, processed and sequenced at scale, then taxonomically assigned with a measure of confidence using bioinformatic methods.^29–31^ When assessing the sustainability of forest management practices, below-ground processes mediated by soil bacteria, fungi, and arthropods should be taken into consideration and these DNA-based tools can help with this.^9^

Wildfire has been shown to affect soil organisms crucial for nutrient cycling, soil stabilization, and remediation, but it can be challenging to study below-ground systems using conventional methods.^9,32,33^ DNA-based methods can help open up this ‘black box’.^34^ In the literature, bacterial communities following wildfire, particularly those associated with charcoal, have been conducted mostly using DNA-based methods such as qPCR, RFLP typing, and more recently high throughput DNA metabarcoding.^35–39^ In studies that survey the microbial community of bacteria and fungi, molecular methods have been useful to facilitate rapid taxonomic identifications with finer resolution. The fungal community following wildfire has been relatively well studied, using conventional above-ground sporocarp sampling, using field or bioassay ectomycorrhizal root tip morphotyping, but also using DNA-based methods.^40–45^ Fire-associated fungal species and the recolonization of surface soils from the fungal spore bank, sclerotia, or surviving mycorrhizae from deeper soil layers have been studied.^41,46–48,49^ Arthropod communities following fire, or following the addition of wood ash, have also been studied through the collection and morphological identification of individuals, but also more recently using metabarcoding.^50–56^ As a result multi-marker DNA metabarcoding is well positioned to further our understanding of community succession and assembly processes in soils, allowing researchers to simultaneously monitor phylogenetically diverse taxa.

When the successional trajectories of interest exceed the lifespan of investigators, sites at different time points since disturbance can be assessed using a chronosequence approach providing a long-term perspective that would otherwise not be possible through repeated measures at one site.^57^ In this study, DNA metabarcoding allowed us to simultaneously detect shifts in the soil microbiome, bacteria and fungi, as well as the soil zoobiome, arthropods along an 80-year chronosequence following stand-replacing wildfires.^27,58^ The objective of this study was to establish a post-fire temporal baseline of soil community recovery in boreal jack pine dominated forests. The results from this study will improve our understanding of changes in the community assembly processes through time. This natural disturbance baseline will provide a point of comparison to human disturbance as a reference condition and can help us assess the importance of older age classes in forests that typically suffer a truncation of age class distribution in managed landscapes due to repeated harvests with shortened timber extraction rotation ages.^59^

## Results

### Establishing a post-fire temporal baseline of soil community recovery

We compared bacterial, fungal, and arthropod communities across fire-origin sites spanning 5 stand development stages (Table 1, Fig 1). Alpha diversity, sample-based richness of sequence variants (SVs), increased from the establishment to crown closure stage (Wilcox test, p-value = 0.01), then plateaued (Fig 2a) for all species groups. Pairwise beta diversity was assessed between successive stand development stages using Jaccard dissimilarity (Fig 2b). To determine the importance of two possible processes driving this dissimilarity, species turnover or species nestedness (loss of species from sites with higher to lower richness), we also present the turnover and nestedness components. We found that the turnover component contributed more towards Jaccard dissimilarity, indicating the relative importance of species replacement from one stand development stage to the next. Gamma diversity, the total SV richness at each stand development stage, is highest for soil bacteria followed by fungi and athropods. For all taxa, gamma diversity peaks at the crown closure stage and is lowest at the establishment stage (Fig 2c).

**Fig 1.**
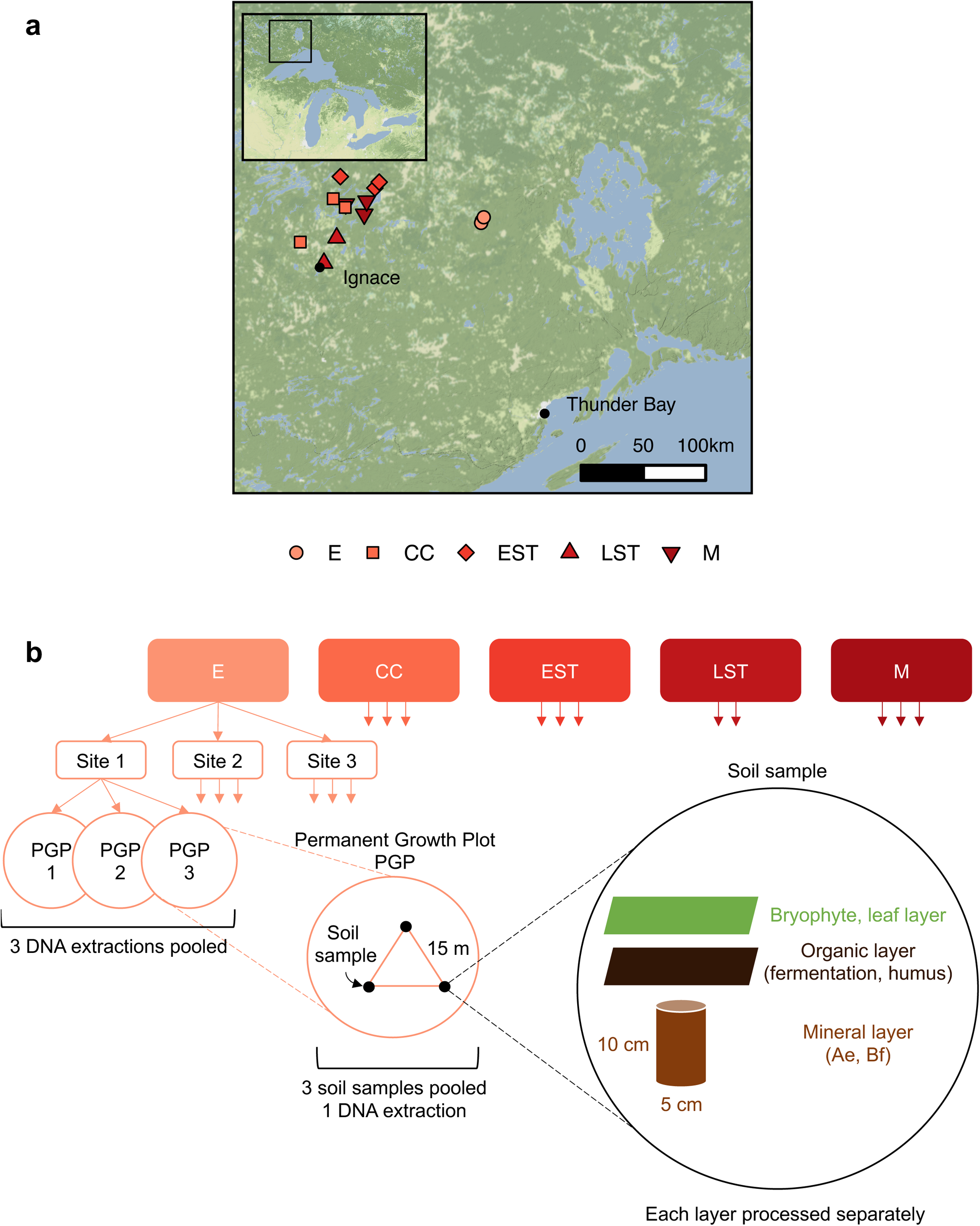
Overview of experimental design. Site locations are shown in part a) Inset map shows the Great Lakes region and the square box indicates the main map area. Scale bar shows 50 km increments. Sampling approach is shown in b) Up to three sites were sampled per stand development stage. Within each permanent growth plot, 3 soil samples were collected and the layers were processed separately. For each soil layer, samples were homogenized and DNA was extracted. DNA extracts were then combined at the site level, again keeping the layers separate. Abbreviations: establishment (E), crown closure (CC), early self-thinning (EST), late self-thinning (LST), mature (M). Map tiles by Stamen Design, under CC BY 3.0. Data by OpenStreetMap, under ODbL.

**Fig 2.**
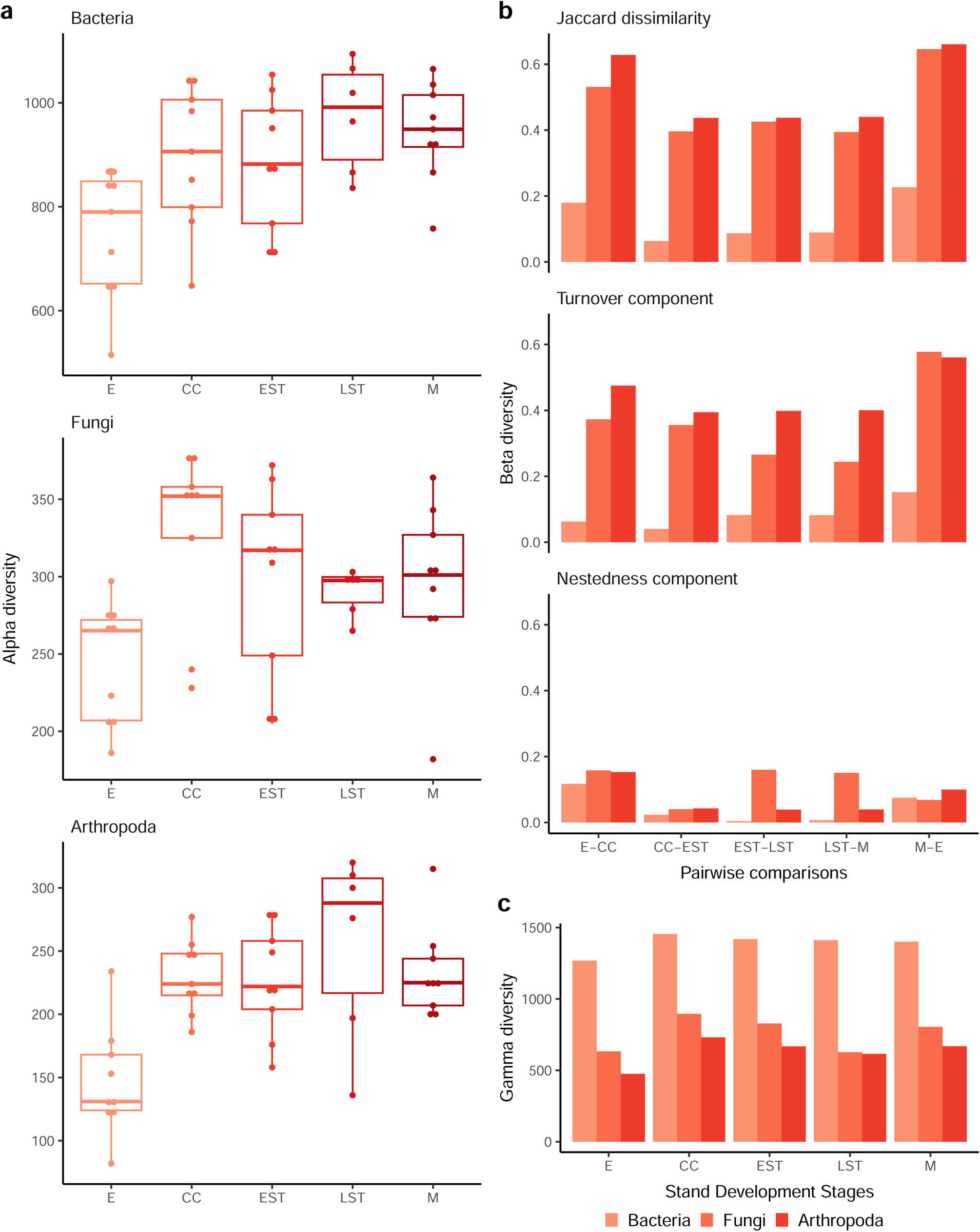
Contrasting patterns of diversity recovery seen for soil bacteria, fungi, and arthropods. Recovery trends across all soil layers are shown in terms of: a) alpha diversity based on sequence variant (SV) richness, b) beta diversity based on Jaccard dissimilarity including the turnover and nestedness components, and c) gamma diversity based on total SV richness across all sites at each stand development stage. Abbreviations: establishment (E), crown closure (CC), early self-thinning (EST), late self-thinning (LST), mature (M).

**Table 1.**
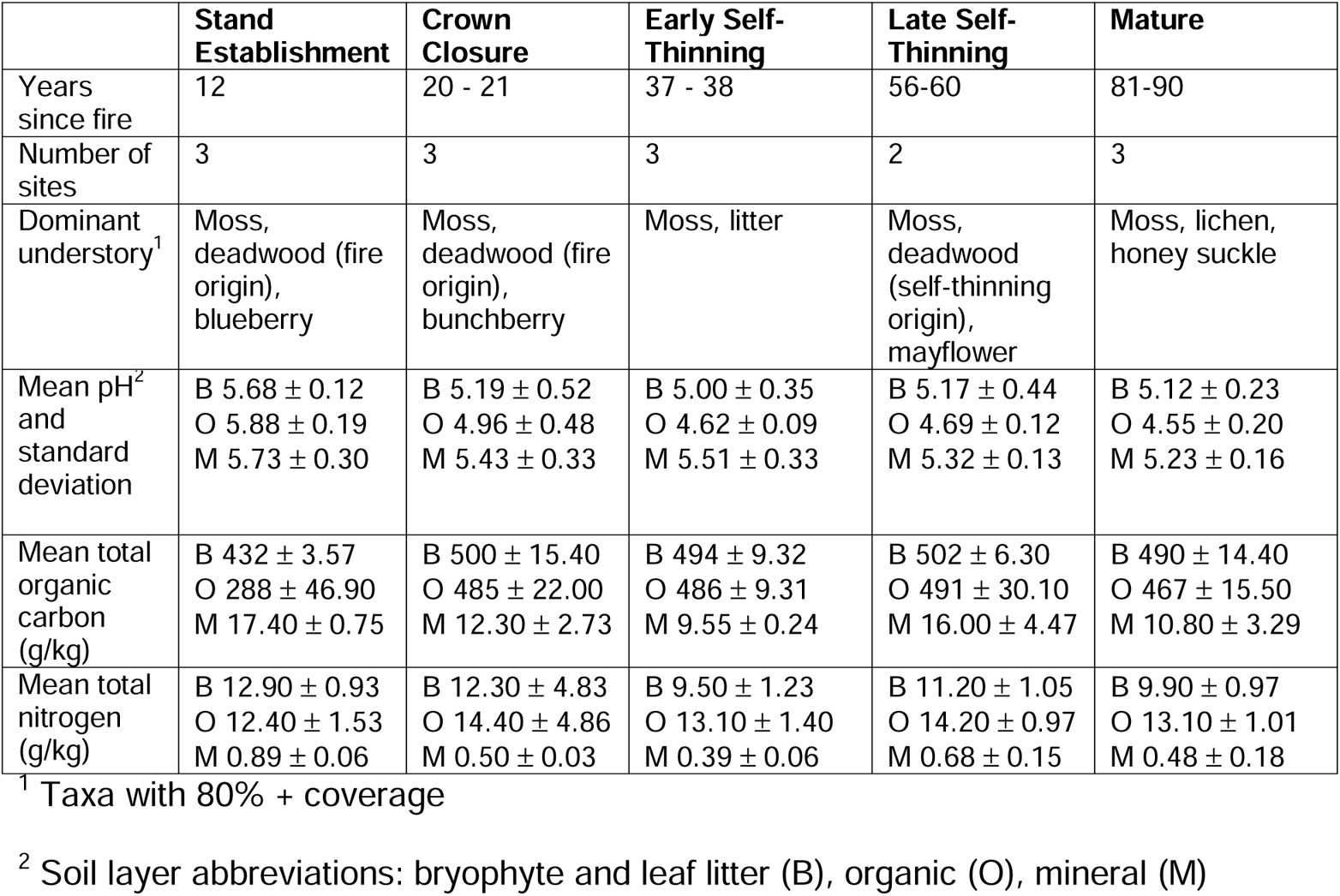
Overview of site characteristics.

Indirect gradient analysis using non-metric multi-dimensional scaling (NMDS) indicated a shift in the species composition across each stand development stage for each organismal group (Fig 3). Both soil bacteria (stress=0.07, linear R^2^= 0.98) and soil arthropods (stress=0.10, R^2^=0.95) were distinct especially at the establishment stage. In contrast, soil fungi were distinct at each stand development stage (stress=0.08, R^2^=0.97). Soil communities at the establishment stage were correlated with higher pH (p-value < 0.05). Subsequent stand development stages were correlated with decreasing pH (p-value < 0.05) and increasing total organic carbon (p-value < 0.05). Variance in beta diversity was measured using Sorensen dissimilarities and was homogenous across successive stand development stages for all taxa. This indicates that differences in community composition are driven largely by differences among the stand development stages. For bacteria, fungi, and arthropods stand development stage was found to explain 44.7%, 40.6%, and 32.2% of the variance in beta diversity (PERMANOVA, p = 0.001).

**Fig 3.**
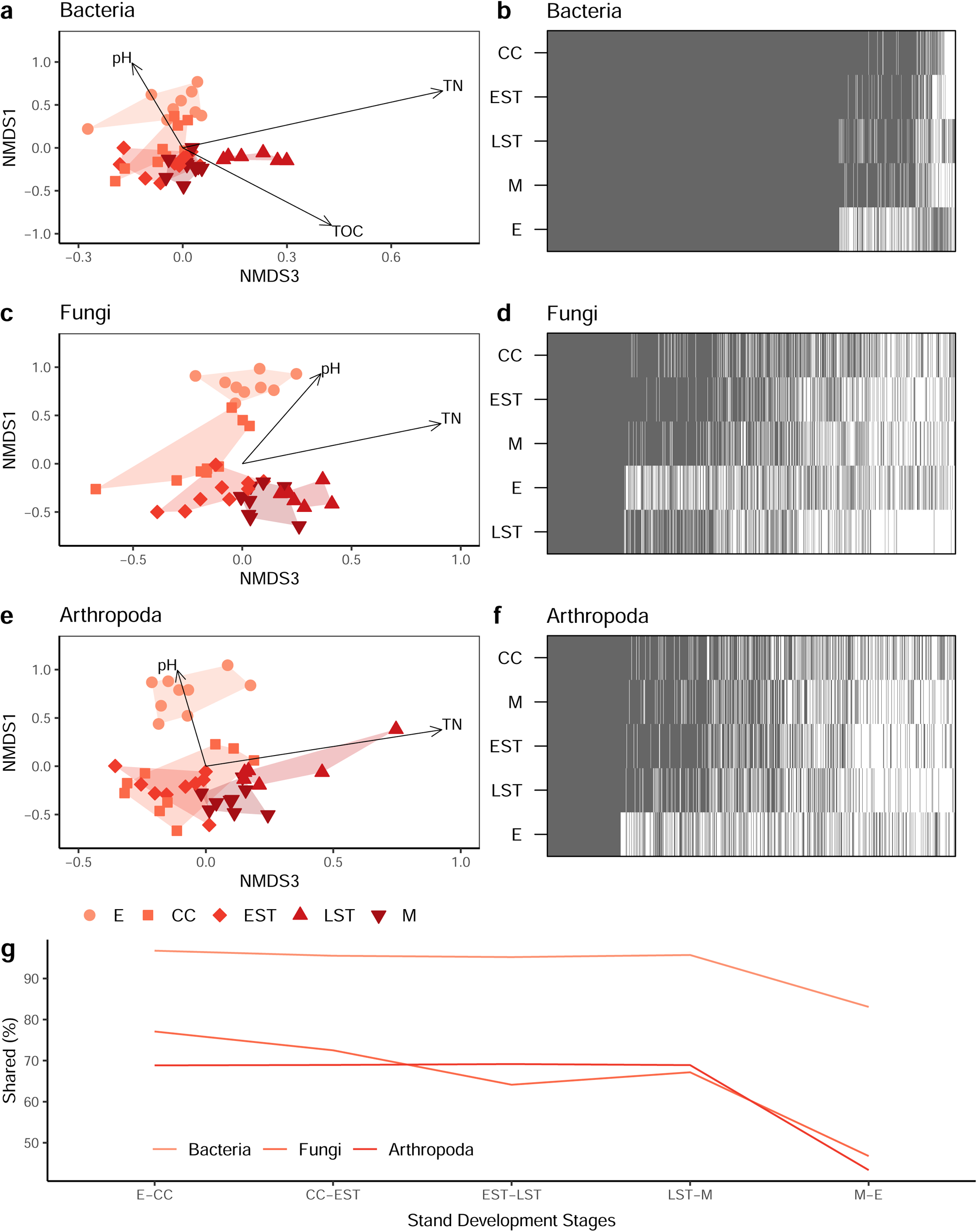
Contrasting recovery patterns of soil taxa along an 80-year chronosequence following natural wildfire disturbance. Community shifts for soil bacteria (a,b), fungi (c,d), and arthropoda (e,f) are shown using non-metric multidimensional scaling plots of Sorensen dissimilarities (left column) and whole community nestedness based on overlap and decreasing fill (NODF) (right column). The percentage of sequence variants shared from one stand development stage to the next is also shown (g). Abbreviations: total nitrogen (g/kg) (TN), total organic carbon (g/kg) (TOC); establishment (E), crown closure (CC), early self-thinning (EST), late self-thinning (LST), mature (M).

Pairwise permutational analysis of variance (PERMANOVA) comparisons across successive stand development stages showed that community composition was usually significantly different between each successive development stage for all taxa (p-value ~ 0.01). The exception was for bacteria, whose community composition was not significantly different between the crown closure and self-thinning stage (p-value > 0.05). Even at broader levels of resolution, i.e., from SVs to genera to family rank, soil communities still appear to be distinct from one development stage to the next (Fig S1). The exceptions are for the bacterial community resolved to the genus rank that is not significantly different from the crown closure to self-thinning stages and the fungal community resolved to the genus rank where there was not a significant difference between the late self-thinning to mature stages.

### Processes that describe community assembly patterns

Whole-community nestedness analysis helped us assess how species were appearing and disappearing from stages over time (Fig 3b,d,f). Nestedness based on overlap and decreasing fill (NODF) measures nestedness both in terms of species compositions (highest richness in top row, lowest richness in bottom row) and in terms of species occupancy across development stages (highest occupancy towards the left, lowest occupancy towards the right). NODF is measured on a scale from 0 to 100 (perfect nestedness) and significance is assessed with respect to a null distribution. The NODF profiles show the proportion of sequence variants that form the core community and are found across every stand development stage (stacked grey blocks towards the left). The grey and white bars to the right represent sequence variants that appear (grey) and disappear (white) across stand development stages.

In contrast with Fig 2b that focused on the drivers of community dissimilarity, Jaccard turnover and nestedness, Fig 3 looks at the whole community nestedness or connectance, species common across all stages and those that come and go. Soil bacterial (NODF 43.8, p-value 0.001), fungal (64.9, 0.001), and arthropods (63.4, 0.001) were significantly more nested than null expectations. Note that in contrast with calculations of nestedness temperature, SVs present in each stand development stage do not contribute towards the NODF score, avoiding over-estimation of nestedness structure. ^60,61^ In Fig 3b,d,f, the order of nestedness provides insights into succession for all taxa: 1) the crown closure stage is always at the top, suggesting that some of the species composition for later stages are inherited from the crown closure stage, 2) establishment communities are nested below mature communities suggesting that at least part of the mature community carries over into the establishment stage following wildfire disturbance.

Bacterial community composition (rows) appears to be chronologically nested from the crown closure stage onwards (Fig 3b). A large proportion of the bacterial community is also similar across all stand development stages (grey block to the left), a result corroborated by low Jaccard dissimilarity values from Fig 2b and clustering of stand development stages in Fig 3a. This result contrasts with fungal and arthropod communities where stand development stage richness increases from the establishment to crown closure stages, then decreases to the early self-thinning and late self-thinning stages, followed by an increase at the mature stage (Fig 3d,f). Generally, fungi and arthropod communities show smaller core communities, with a smaller subset of species being inherited through successional stages following crown closure. Each stage harbours its own unique community of successional species that shift over time, a result corroborated by the ordinations shown in Fig 3c,e. We also calculated the proportion of sequence variants shared between pairwise comparisons of successive stand development stages. For example, when comparing the establishment to crown closure stage (E-CC), 97% of bacterial sequence variants from the establishment stage were also detected in the crown closure stage. Approximately 95-97% of bacterial and 64-77% of fungal and arthropod sequence variants were shared between stages. Following wildfire disturbance, about 83% of bacteria and 36-43% of fungi and arthropod sequence variants from the mature stage were detected in the establishment soil communities.

### Simulating the truncation of stand development stages

We simulated shorter fire return intervals and losing older age classes (Fig 4) on gamma diversity, the total number of unique sequence variants across all sites, and found that gamma diversity decreased as later stand development stages are lost, especially for fungi and arthropods (Fig 4b). This implies that each stand development stage is important for maintaining total soil biodiversity, especially for fungi and arthropods. We also found that whole community nestedness decreased for all taxa, and again, especially for fungi and arthropods (Fig 4c). Nestedness was significantly greater (p-value < 0.05) than null expectations when at least three of the early stand development stages are present. When only establishment and crown closure stages are compared, nestedness is not significantly different from a randomized community. These results suggest that communities may be becoming less stable and more susceptible to the loss of rare species as later stand development stages are lost. We also assessed nestedness and specialization patterns for the major fungi taxonomic and functional groups. Fungal undefined saprotrophs, ectomycorrhizal fungi, and all major taxonomic groups showed greater nestedness than null expectations (p-value < 0.05) (Fig S2a, S3). All major fungal groups also showed greater specialization to a particular stand development stage than null expectations (p-value = 0.001)(Fig S2b). This suggests that stand development stage may ‘select for’ particular fungal taxa and functions, a result corroborated by the large number of stand specific bioindicators that were detected in this study, described in the next section.

**Fig 4.**
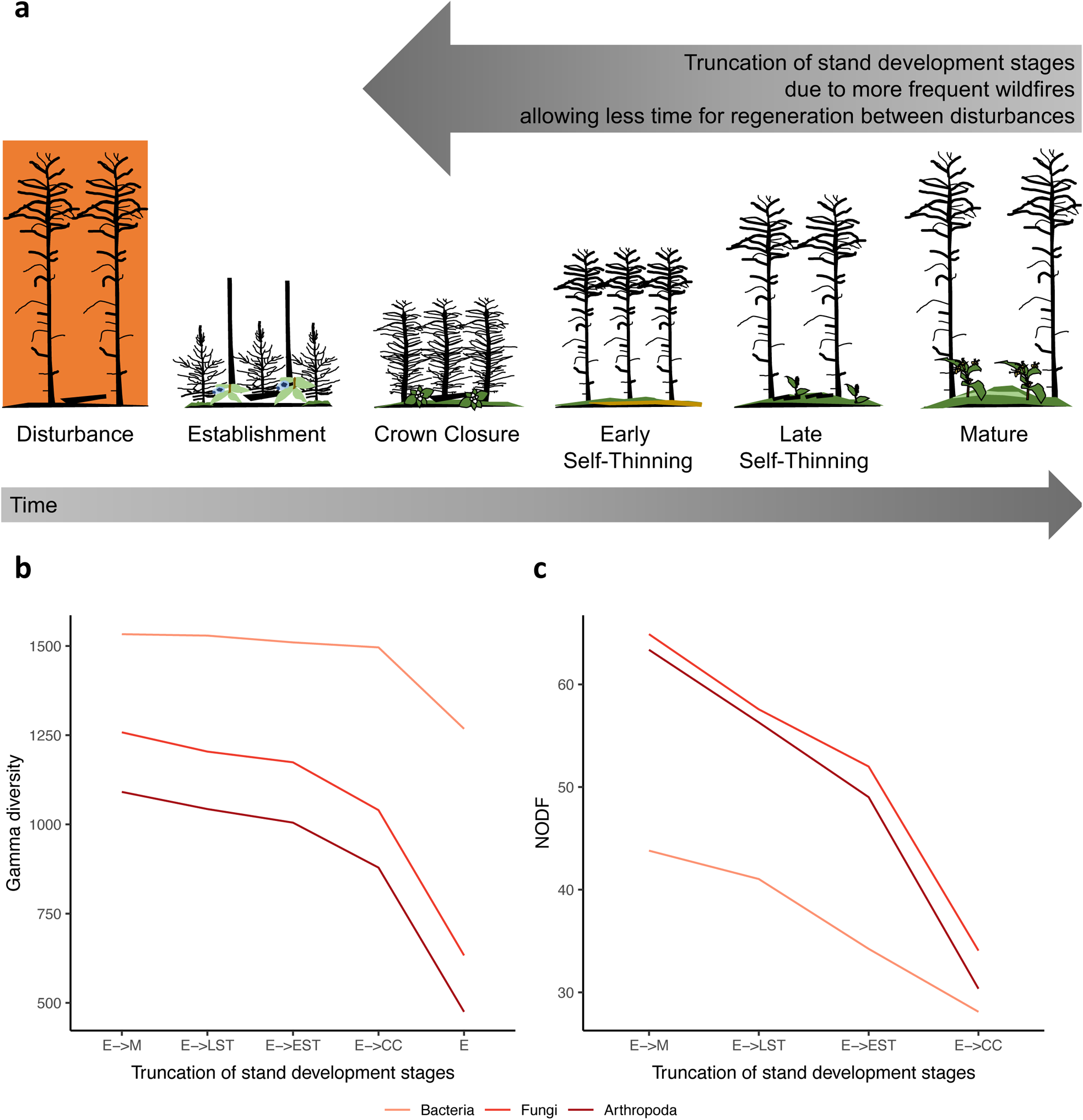
Each stand development stage is important for maintaining gamma diversity and whole community nestedness. Stand development stages for a) the full cycle, establishment to mature (E->M), is compared with truncated cycles where the oldest age class is sequentially removed. We show changes in simulated b) gamma diversity and c) nestedness based on overlap and decreasing fill (NODF). For NODF, nestedness is statistically greater than null expectations (p-value < 0.05) for each cycle except for the shortest (E->CC). Abbreviations: establishment (E), crown closure (CC), early self-thinning (EST), late self-thinning (LST), mature (M), nestedness based on overlap and decreasing fill (NODF).

### Identifying taxa and functions associated with post-fire recovery

The core communities, taxa detected at each stand development stage, were also described. The major fungal taxonomic groups detected across each stand development stage included the Agaricomycetes (many produce mushrooms), Leotiomycetes (many are plant pathogens), Dothideomycetes (diverse ecological roles), Eurotiomycetes (diverse ecological roles, some produce toxic or useful secondary metabolites), and unidentified fungi (Fig 5a). For the fungal community, we also mapped ecological function to assess the trajectory of recovery for major functional groups (Fig 5b). It can be informative to compare changes in diversity based on taxonomic groups, as well as based on function, decoupled from taxonomy. The abundance of major taxonomic and functional groups varied across each stand development stage, with two notable trends: 1) a decrease in the relative abundance of fire-associated fungi and 2) an increase in the relative abundance of ectomycorrhizal fungi from the establishment to onset of self-thinning stage.

**Fig 5.**
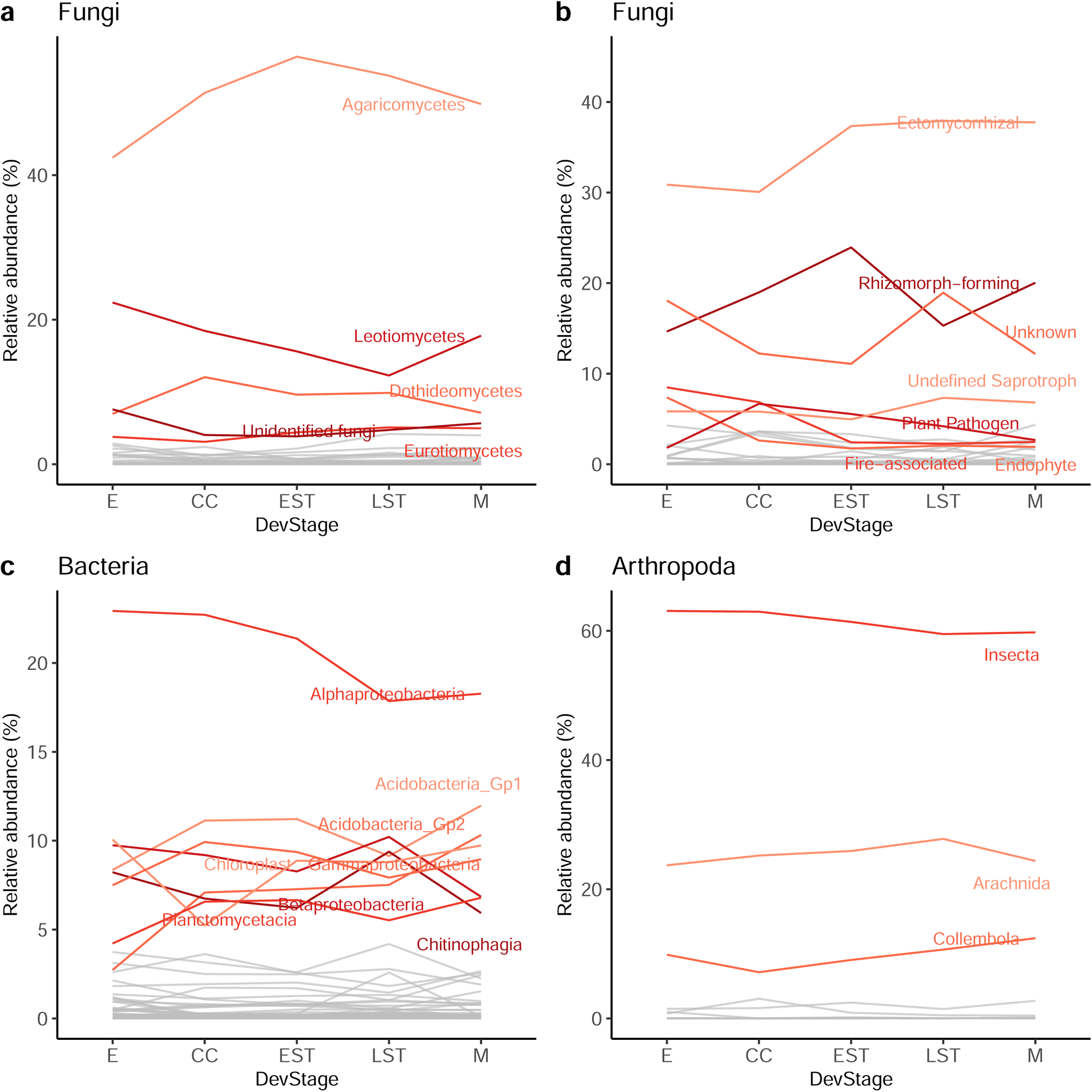
Major groups vary across the post-fire chronosequence. The relative read abundance of a) fungal classes, b) fungal ecological guilds and traits, c) bacterial classes, and d) arthropod classes are shown. Major groups that comprise at least 5% of the relative read abundance in at least one stand development stage are highlighted in color and labelled. Abbreviations: establishment (E), crown closure (CC), early self-thinning (EST), late self-thinning (LST), mature (M).

We expected that fire-associated taxa would be found at earlier stand development stages based on the existing literature. To ensure that DNA metabarcoding was successfully detecting taxa that we would expect to be present in our sites, we screened our samples for pyrophilous bacteria, fungi, and arthropods. For fungi, 8 species of previously known fire-associated fungal taxa, based on conventional survey methods in the literature, were detected in this study including *Anthracobia macrocystis*, *Coltricia perennis*, *Coprinellus micaceus*, *Gymnopilus decipiens*, *Hygrocybe conica*, *Morchella exuberans*, *Sphaerosporella brunnea*, and *Thelephora terrestris* (Fig S4, Table S1). Because 46 out of 105 (43%) fire-associated fungal species from previous studies were not represented in our reference sequence database, we also summarized our results to the genus rank where only 8 out of 56 (14%) targeted fire-associated fungal genera were missing. Twenty-four additional taxa that are known to be associated with fire were also detected. Of these, unidentified Pyronemataceae were represented by 4 unique species hypothesis (SH) groupings. In the UNITE database, these SHs were comprised of sequences belonging to the heat-resistant *Trichophaea abundans*, the cosmopolitain post-fire *Anthracobia melaloma*, as well as the fire-associated cup-fungi genera *Scutellinia,* and *Byssonectria*.^32,62–64^ The frequency and diversity of fire-associated fungal species was highest at the establishment stage, declining over time. Fungal genera, on the other hand, appeared cosmopolitan across development stages and likely includes species that are not strictly fire-associated.

Bacterial communities were dominated by bacterial classes expected to be present in forest soils such as Alphaproteobacteria, Acidobacteria, Gammaproteobacteria, Betaproteobacteria, Planctomycetacia, and Chitinophagia (Fig 5c).^65^ Many bacterial putatively fire-associated genera were found across all stand development stages (Fig S5). An assessment of bacterial functional profiles, however, was difficult as ~ 60% of bacterial reads could not be assigned and the remaining could only be assigned high-level functional classifications such as chemoheterotrophy, aerobic chemoheterotrophy, and chloroplast functions.

The dominant arthropod taxa included Insecta, Arachnida, and Collembola (Fig 5d). Many arthropod putatively fire-associated genera were found across all stand development stages (Fig S6). An assessment of arthropod functional profiles was also challenging because online databases often focus on traits for freshwater rather than terrestrial taxa. Although we were able to create a reference set of traits compiled based a combination of online databases and literature searches, up to ~25% of arthropod reads could not be functionally assigned. Additionally, some arthropod families represent such a diverse array of ecological functions that functional assignment to the species rank would be ideal but not feasible using the semi-automated methods applied in this study.

We also looked for additional SVs that were significantly correlated with each development stage (Table S2). The number of stand development stage bioindicators in fire-origin sites was greatest for bacteria (381 indicator SVs/1533 SVs total, 24.8% bioindicator SVs), followed by fungi (296/1258 SVs, 23.5%), and least for arthropods (200/1091, 18.3%). The number of bioindicator SVs was highest for the establishment stage (357 in total, 180 bacteria, 128 fungi, 49 arthropods). Taxa associated with the establishment stage include bacterial *Sphingomonas*, *Mucilaginibacter*, and *Streptophyta*; fungal *Suillus* and *Pezoloma*; and arthropod *Microcerotemes* and *Phortica*. The number of bioindicators was lowest in the crown closure and early self-thinning stages (93 and 65 SVs respectively). The second highest number of bioindicators were for the late self-thinning stage (242 in total, 102 bacteria, 73 fungi, 67 arthropods). Taxa associated with the late self-thinning stage included bacterial *Mucilaginibacter*, *Flavitalea*, and *Phenylobacterium*; fungal *Piloderma* and *Tomentella*; and arthropod *Moritzoppia*. The third highest number of bioindicators were for the mature stage (120 in total, 61 bacteria, 26 fungi, 33 arthropods). Taxa associated with the mature stage included bacterial *Granulicella*, *Acidibacter*, and *Roseiarcus*; fungal *Archaeorhizomycetes* and *Infundichalara*; and arthropod *Ceratozetes*, *Entomobrya*, *Moritzoppia*, *Orconectes*, *Winthemia*, and *Zhuginia*. These additional results further suggest that stand development stages are distinct communities that are significantly correlated with specific sets of taxa.

## Discussion

Our study demonstrates that each successional stage contributes unique biodiversity in multiple taxonomic groups. Of the groups examined here, this uniqueness is more evident for fungi and soil arthropods than for bacteria which appear to be more ubiquitous. This result has significant implications for the future of soil biodiversity in a changing climate where fire return intervals are expected to shorten, resulting in a truncation in the age class distribution across broad ecoregions.^66^ More specifically, habitats for the biodiversity unique to the oldest forests are expected to decline. From a forest management perspective within the managed forest landscape, our results emphasize a recognition within the current forest management planning and harvest scheduling approach (i.e., currently the “oldest first” approach) that will ensure adequate traits of the old forests (e.g., old growth) are being maintained.

Fire-associated bacteria, fungi, and arthropods do not necessarily require fire to complete their life cycles, but may share traits such as heat resistance, are cosmopolitan, have thermotolerant structures, fast colonization/growth, and the ability to acquire resources in an environment altered by fire.^37,45,51,64,67–69^ For instance, pyrophilous or fire-loving fungi often appear after fire and represent a range of species that may be fire-adapted, cosmopolitan, or opportunists; some pyrophilous fungi have even been identified living as endophytes inside mosses and living as endolichenic fungi of lichens.^45–47,64^ For example, *T. terrestris*, known to disperse quickly and colonize roots from spores, was found more frequently in our study in early stages, but was detectable from every stage.^70^ This is consistent with a previous study in a jack pine dominated chronosequence that found *T. terrestris* to be one of the dominant ectomycorrhizal fungi associated with early stage samples 5-11 years post-wildfire.^70^ Here, DNA metabarcoding detected known pyrophilous fungi that were more diverse and frequent in establishment sites compared with later stand development stages as well as fungal SVs significantly correlated with post-fire sites at the establishment stage.

An unexpected taxon detected in our study, Archaeorhizomycete fungi, does not form above-ground sporocarps so are not typically known as a pyrophilous taxon, however, recent studies that have used DNA metabarcoding have found these fungi to be abundant following wildfire.^38,71^ Archaeorhizomycete fungi were initially described from marker gene studies of soil, found to have a global distribution, and later two species were isolated as slow-growing cultures.^72–74^ In a study of soil fungi from a subtropical peatland, Archaeorhizomycete fungi had high relative abundance responding to low- and high-intensity fire at various soil depths.^71^ Another study has specifically linked Archaeorhizomycete fungi with decreased soil water repellency and increased soil moisture content, following wildfire in a European pine stand.^38^ In this study, we detected Archaeorhizomyces SVs as bioindicators of establishment and mature stand development stages, representing the 11^th^ most abundant class of fungi based on relative read abundance.

There were, however, pyrophilous taxa that we did not detect in our study. For example, we did not detect *Pyronema* spp. or *Pholiota carbonaria* that have been found to produce fruiting bodies following wildfire.^43,75,76^ We also did not detect any pyrophilous insects. The lack of detection may have been due to a lack of representation in the underlying reference sequence database used making it problematic for identifying pyrophilous insects. Secondly, our first set of samples were at the establishment stage 12 years post-fire whereas most studies of pyrophilous taxa usually occurs within days to weeks following the fire disturbance, so the opportunity to detect these taxa in our study may have passed.

Soil bacteria had a large core community that shared many sequence variants across stages. Community composition was also chronologically nested from the crown closure stage onwards. We saw distinct species-inheritance structures through time, i.e., organisms that colonized earlier persist across stages presumably so long as they have a niche, with other taxa coming and going. These results suggest that soil bacteria may be resistant to fire disturbance and recovery relatively quickly. These findings are in agreement with a meta-analysis of 131 studies that also found that soil bacteria tended to be more resilient to fire than soil fungi, showing a lower effect of fire on bacterial richness and diversity compared with fungi.^42^ Our results are also in alignment with a previous study in a Chinese boreal forest that found that the bacterial community recovered quickly in terms of richness and composition by 11 years, with recovery correlated with edaphic soil properties such as changes in pH, moisture, and nitrate.^77^ Our natural disturbance results are also consistent with controlled soil heating experiments that showed relatively rapid bacterial recovery compared to fungi, where the authors suggested that higher soil pH favored bacterial growth and possible competitive interactions between soil bacteria and fungi that resulted in extended recovery time for fungi.^78^ When describing recovery following wildfire, bacteria may be better suited for assessing early successional changes rather than for assessing longer term changes beyond the crown closure stage.

Soil fungi and arthropods in this study shared relatively small core communities carried through each stand development stage and each stage appeared to support distinct and unique community assemblages. For these groups, the order of nestedness in terms of species composition was not strongly related to chronology, rather there is an increase in richness from the establishment to crown closure stage, then there is species loss with the onset of self-thinning and late self-thinning stages which were compositionally nested within the crown closure stage, followed by an increase in richness at the mature stage. Compared with bacteria, a greater proportion of fungal and arthropod sequence variants come and go suggesting dynamic reshaping of the communities that become more or less diverse from one stage to the next following crown closure. These results are consistent with a previous study that found ectomycorrhizal succession after fire was best described with respect to stand development stage as opposed to categories such as early/multi-staged/late successional types.^79^ The large number of stand development stage bioindicator taxa we detected further supports this stand development stage approach to post-disturbance assessment of biotic recovery. To maintain the full suite of biodiversity in jack-pine dominated boreal forest soil communities, it will be important to maintain a mosaic of stand development stages across broad forested landscapes.

Increases in the frequency, intensity, and scale of wildfires in boreal regions have been predicted due to climate change. ^1,2,66,80–82^ In addition, there is evidence that current forest harvest levels over large scales has also resulted in a truncation of age class distributions which is also likely to result in a reduction in biodiversity associated with these older age classes.^59^ In this study, we show a simulation where, as recovery time decreases (i.e., truncation of later stand development stages), we see decreasing landscape-level diversity. Although simple diversity metrics like richness appear to plateau quickly, the underlying community composition can take significantly longer to ‘recover’, especially for fungi and arthropods. In jurisdictions where forest management policy, guidelines, and practices are mandated to emulate natural disturbance, simply looking for convergence across stand development stages may not be overly effective when assessing sustainability since this approach may not adequately account for changes in landscape-scale diversity. Successional species adapted to specific stand development stages, especially later stages, could be lost if not actively managed for. Forest management policies, planning, and practices should not only continue to actively control/suppress wildfires but should also be designed to maintain a mosaic of stand development stages across broad landscapes units (e.g., FMU – forest management unit and/or ecoregional scales) to ensure the taxonomic and functional diversity they collectively represent are being conserved.

Another consequence of truncated recovery times following wildfire disturbance, is the decreasing stability of soil communities. In this study, when we simulated the loss of older age classes, we observed a decrease in whole community nestedness. Based on the network interaction literature, increased nestedness or connectedness leads to increased robustness; whereas decreased connectedness may result in communities that are more susceptible to random extinctions.^61,83^ In a jack pine system that depends on ectomycorrhizal fungi, an interruption or truncation in community assembly processes could hinder forest recovery following wildfire.^84^ Although in the literature fire has been described as a driver of fungal diversity^45^, wildfire disturbance initially decreases the diversity of all taxa in the short term especially when fire severity is high.^77,85^ Fungal taxa and functional groups have also been shown to vary with respect to fire severity tolerance with basidiomycetes and ectomycorrhizal fungi shown to have relatively low tolerance to fire.^86^ In our study, the relative abundance of ectomycorrhizal fungi did not plateau until the onset of self-thinning stage (20 – 37 years) yet the underlying ectomycorrhizal community composition was distinct at each stand development stage. As a comparison, in a British Columbia mixed temperate forest ectomycorrhizal fungal diversity plateaued by 26 years but community composition did not stabilize until 65 years.^79^ If wildfire frequency continues to increase under climate change as predicted, the recovery time of important ecological guilds such as ectomycorrhizal fungi could be short-circuited, ultimately affecting nutrient dynamics in boreal forest soils, and, in turn, may effect tree seedling establishment and growth.

## Conclusions

Soil DNA metabarcoding was an effective tool to simultaneously survey a comprehensive set of taxa across diverse groups of bacteria, fungi, and arthropods. This study contributes to the literature by broadening the window of observation of the occurrence and distribution of pyrophilous taxa, but future studies would benefit from improved reference sequence databases to minimize false-negatives (false-absences). The utility of this technique should improve over time with continued growth in taxonomic and functional reference databases. Community differences across stand development stages had a large turnover component, implicating the importance of species replacement during succession. Our results emphasize the need for a forest management strategy that actively represents the complete range of stand development stages to maintain a full suite of biodiversity, including successional species, especially for fungi and arthropods. The biotic characterization of soils in these development stages will act as an important baseline for future work to assess the stability of soil networks and sustainability of harvest. Lastly, silvicultural practices such as vegetation control may alter the natural successional pathways and have the potential to change soil faunal assemblages. To examine this explicitly we are planning future chronosequence work to examine the post-harvest biodiversity relative to the post-fire biodiversity to better understand the implications of harvest disturbance on soil biodiversity over the long-term through all successional stages.

## Methods

### Study Site Selection

This study used the chronosequence approach that replaces time (repeated measures) with space, with the underlying assumption that the primary differentiating factor between sites is time since disturbance.^87^ To conform to the assumption noted above, our principal selection criterion was based on similarity in the basic site characteristics (*e.g.*, climate, soils, tree species composition, and productivity). Sites were separated by at least 1 km and represent independent stands, although some were the result of the same large-scale wildfire events. All of the sites experienced severe fires with Fire Weather Index (FWI) values ranging from 17.8 to 44.6. A total of 14 sites were included in this study (Table 1, Fig 1) from wildfire origin stands. These sites were selected to represent stand development stages, with up to three replicates per stand development stage. Beyond confirmed stand age, final selection and placement within stand development stages was based on key stand attributes (i.e., establishment - uniformly established pine regeneration along with low shrub and herbaceous species present; crown closure - interlocking crowns with some evidence of self-pruning and a reduction of herbaceous cover; early self-thinning - some canopy differentiation, with smaller trees beginning to die; late self-thinning - most lower canopy pine dead, with spaced out larger trees; mature - some random mortality of dominant trees, with some ingressed black spruce in the lower to mid-canopy).

All study sites experience cold continental climates, with annual precipitation ranging from 68 to 82 cm yr^−1^ (growing season precipitation from 41 to 49 cm yr^−1^), annual mean temperatures of 1.0°C to 1.8°C, and Growing Degree-Days (GDD) from 1112 to 1263. The soils for all of the study sites represent well-drained, outwash sands, with well-developed Orthic Humo-Ferric Podzolic profiles. All stands were jack pine-dominated (*Pinus banksiana*), with many having minor amounts of black spruce (*Picea mariana*), trembling aspen (*Populus tremuloides*), and white birch (*Betula paperifera*). The majority of the stands are classified as Site Class I (Plonski 1981), with estimated site index values ranging between 18-22 m at breast height age 50.

### Soil Sampling and Laboratory Analysis

As part of a larger project, 3 – 400 m^2^ (11.28 m radius) permanent growth plots (PGPs) were randomly established at each study site and were located at least 50 m apart from each other. Three 15 m transects (9 per site) to measure downed woody debris were superimposed over each of the PGPs. The first transect was oriented based on a random azimuth, with the remaining two transects offset 120° to form a triangle. Soil sampling was done at each transect endpoint (points of the triangle), giving 9 soil sampling locations per site.

In 2017, soil profiles at each of the 9 within-site locations were sampled. A 5 cm x 10 cm x full depth volume of the bryophyte and/or leaf layer was sampled. The same volume of the organic horizon (i.e., LFH: litter, fermentation, and humus layers) was also sampled. Upper mineral soil (MIN, 0-10 cm) was collected separately. Soil was extracted using a 5 cm diameter PVC pipe driven to a depth of 10 cm into the mineral soil. PVC pipes were sanitized with bleach or concentrated alcohol before sampling was performed. All sampling was completed with gloves to reduce cross-contamination. Samples were immediately frozen at −20°C after sampling and held at that temperature until analysis. The collected mineral soil consisted of a mix of the Ae and upper Bf horizons, which, through pedogenetic processes, have different physical and chemical properties. Additional samples of the LFH and mineral soil horizons (Ae, Bf collected separately) were also collected for pH and chemical determinations. Once back in the laboratory, fresh (field moist) subsamples were tested using an Oakton pHTestr5 for distilled water and CaCl_2_ pH in a 2:1 slurry that was shaken and let stand for 10 minutes before the pH reading was taken. The remaining samples were air-dried, and either ground in a Wiley mill (forest floor samples) or sieved through a 2mm sieve and ground using a mortar and pestle (mineral soil samples). Total soil C concentrations (%) was determined by dry combustion using a LECO CNS-2000 analyzer (LECO Corporation, St. Joseph, MI, USA). Soil nitrogen (N) concentrations were determined by the semi-micro-Kjeldahl procedure (Kalra and Maynard 1991). Exchangeable cations were determined by ICAP in unbuffered 1 mol L^−1^ NH_4_Cl solution.^88^ Extractable phosphorus (P) was determined by ICAP in Bray and Kurtz No. 1 extractant.^88^ Individual concentrations were adjusted for air-dried moisture content and reported on an oven-dried basis (mg kg^−1^).

### Molecular biology methods & sequencing

The samples from each PGP were combined but the layers were kept separate. Bryophyte/litter and organic layers were each separately homogenized in a knife mill and mineral soil was homogenized by forcing it through a 0.2 mm sieve. The knife mill and sieve were rinsed with water and cleaned with 70% ethanol between samples. DNA extractions performed by NRCan lab using 0.25 g of soil with the MoBio PowerSoil DNA Isolation Kit following the manufacturer’s instructions except that 200 ul of 100 mM AlNH_4_(SO_4_)_2_ was added to the tube along with soil and Solution C1 followed by a 10 minute incubation at 70°C to help lyse cells.^89^ DNA extracts were then combined at the site level, resulting in one combined DNA extract per site per layer. Bacterial and fungal communities were enriched by targeting the 16S v4-v5 region using the 515F-Y (5’ - adapter + *primer* - 3’) TCGTCGGCAGCGTCAGATGTGTATAAGAGACAG + *GTGYCAGCMGCCGCGGTAA* and 926R GTCTCGTGGGCTCGGAGATGTGTATAAGAGACAG + *CCGYCAATTYMTTTRAGTTT* primers; and the ITS2 region using the ITS9 TCGTCGGCAGCGTCAGATGTGTATAAGAGACAG + *GAACGCAGCRAAIIGYGA* and ITS4 GTCTCGTGGGCTCGGAGATGTGTATAAGAGACAG + *TCCTCCGCTTATTGATATGC* primers. ^90–92^ Invertebrate communities were enriched using two sets of primers targeting the COI gene F230R_modN marker using the LCO1490 (5’ - adapter + *primer* - 3’) TCGTCGGCAGCGTCAGATGTGTATAAGAGACAG + *GGTCAACAAATCATAAAGATATTGG* and the 230R_modN GTCTCGTGGGCTCGGAGATGTGTATAAGAGACAG + *CTTATRTTRTTTATNCGNGGRAANGC* primer adapted from Gibson *et al*. 2015 to include N’s instead of inosine bases; and the BE marker using the B TCGTCGGCAGCGTCAGATGTGTATAAGAGACAG + *CCIGAYATRGCITTYCCICG* and E GTCTCGTGGGCTCGGAGATGTGTATAAGAGACAG + *GTRATIGCICCIGCIARIAC* primers.^93–95^ For each sample a PCR cocktail of 5 µl template DNA (5 ng/µl in 10 mM Tris pH 8.5), 0.5 µl each of 10 µM forward and reverse primer, 25 µl HotStar Taq plus master mix kit 2x, and 19 µl of sterile water. PCR cycling conditions were as follows: 95°C for 5 minutes followed by 30 cycles (16S) or 35 cycles (ITS) or 40 cycles (COI) of 95°C for 30 seconds, 50°C for 30 seconds, and 72°C for 1 minute, followed by a final extension of 72°C for 5 minutes (10 minutes for COI) and hold at 4°C.

PCR products were visualized on GelRed-stained 1% agarose gels using the ChemiGenius Bioimaging System (Syngene, Cambridge, UK). PCR products were purified using 81 μl of magnetic bead solution (Agencourt AMPure XP, Beckman Coulter Life Science, Indianapolis, IN, USA) according to Illumina’s protocol.^96^ Indexes were added to each sample by amplifying 5 μl of the purified PCR product with 25 μl of KAPA HIFI HotStart Ready Mix, 5 μl of each Nextera XT Index Primer (Illumina Inc., San Diego, CA, USA) and 10 μl of UltraPure DNase/Rnase-Free Distilled Water for a total volume of 50 μl. Thermal cycling conditions were as follows: 3 min at 98°C, 8 cycles of 30 sec at 98°C, 30 sec at 55°C, 30 sec at 72°C, and a final elongation step of 5 min at 72 °C. Indexed amplicons were purified with the magnetic beads as previously described, quantified using a Qubit dsDNA BR Assay Kit (Life Technologies) and combined at equimolar concentration. Paired-end sequencing (2 × 250 bp) of the pools was carried out on an Illumina MiSeq at the National Research Council Canada, Saskatoon. 15% PhiX was added to help compensate for low sequence heterogeneity on the plate.

### Bioinformatic methods

Demultiplexed, paired-end Illumina reads were analyzed on the General Purpose Science Cluster, a high performance computing cluster provided by Shared Services Canada. Metabarcodes were processed using the MetaWorks v1.9.3 multi-marker metabarcode pipeline using the standard exact sequence variant (ESV) workflow.^97^ Briefly, reads were paired using SEQPREP v1.3.2 using the default parameters except that the Phred score quality cutoff was set to 20 and the minimum overlap was set to 25 bp.^98^ Primers were trimmed using CUTADAPT v3.2 using the default parameters except that the minimum sequence length was set to 150 bp, Phred quality score cutoffs at the ends was set to 20, and the maximum number of N’s was set to 3.^99^ Reads were dereplicated and denoised using VSEARCH v2.15.2, setting the minimum cluster size to retain to 3, so clusters with only 1 or 2 reads were removed as noise.^100^ This workflow generates sequence variants (SVs), also referred to as amplicon sequence variants (ASVs), exact sequence variants (ESVs), or zero-radius OTUs (ZOTUs) in the literature, and are clusters of sequences that represent a unique sequence after denoising.^101,102^ The 16S metabarcodes were taxonomically assigned using the RDP Classifier v2.13 using the built-in 16S reference set.^29^ The minimum bootstrap support value was set to 0.80 at each taxonomic rank as recommended. The ITS metabarcodes were taxonomically assigned using the RDP Classifier with a custom-trained reference set based on the UNITE v8.2, QIIME release for Fungi v 04.02.2020, available from https://github.com/terrimporter/UNITE_ITSClassifier.103 Outgroup taxa (single-celled eukaryotes) from the RDP Classifier 2014 UNITE ITS reference were added to the newer custom-trained ITS classifier used here.^104^ A selection of outgroup taxa (plants) were also added from the PLANiTS plant ITS reference set by clustering their dataset by 50% sequence similarity in VSEARCH, and adding the centroid sequence to the ITS classifier used here.^105^ Leave-one-sequence-out testing showed that fungal ITS taxonomic assignments using at least 200 bp queries are at least 90% correct at the kingdom rank and 80% correct at the phylum - family rank assuming that the query sequence is represented in the reference set. Genus level assignments are at least 80% correct when using a bootstrap support cutoff of 0.70. The COI metabarcodes were taxonomically assigned using the RDP classifier with a custom-trained reference set.^31^ As a part of a larger study, we analyzed a total of 4,454,896 reads (16S 1,548,803; ITS 1,540,489; BE 68,396; F230 1,297,208) corresponding to 64,350 unique SVs (16S 45,354; ITS 13,248; BE 992; F230 4,756) from wildfire sites.

### Data analysis

Data was analyzed in Rstudio 2021.09.0 build351 using R v4.1.1.^106,107^ Diversity analyses were carried out with the vegan package.^108^ For each sample, sequence reads were rarefied using the ‘rarecurve’ function to ensure that sequencing depth was sufficient across samples (Fig S7). In subsequent richness and beta diversity analyses, rare SVs that comprised less than 0.01% of reads were further removed as these may be unreliable representing artefactual sequences or affected by tag-jumping.^109^ SV richness was assessed using the ‘specnumber’ function. Normality was assessed visually using the ‘ggqqplot’ function from the ggpubr library and the base R Shapiro-Wilk test ‘shapiro.test’ function, and t-tests were used to compare samples against the reference condition.^15^ “Holm” adjusted p-values are shown for multiple comparisons. Beta diversity was assessed from binary stand development stage x SV matrices to calculate Jaccard dissimilarity as well as the turnover and nestedness components using the nestedbetajac function in the vegan package.^111^ Gamma diversity was calculated as the total number of unique SVs found across all sites and development stages. Differences in community composition were assessed using binary Bray Curtis dissimilarity matrices created using the ‘vegdist’ function with binary=TRUE in PERMANOVA and pairwise PERMANOVA comparisons.^108^ Community shifts were visualized using non-metric multidimensional scaling plots calculated using the ‘metaMDS’ function using a presence-absence matrix and Bray Curtis dissimilarity (Sorensen dissimilarities), using 3 dimensions after running ‘dimcheckNMDS’ from the goeveg package and setting trymax=100.^112^ Stress was assessed using the ‘stressplot’ function. Dispersion was assessed using the ‘betadisper’ function and the base R ‘anova’ function. Environmental variables were fit to the ordination using the ‘envfit’ function with 999 permutations, and only variables with a p-value < 0.05 were plotted. All plots were created using ggplot2 and the map was created using ggmap.^113,114^

Bacterial taxonomic assignments were retained if they had at a bootstrap value of least 0.80 as recommended on the RDP Classifier website.^29^ To assess functional composition, 16S sequences were assigned using FAPROTAX v1.2.4. Since FAPROTAX is optimized to work with SILVA 128 or 132 taxonomies, our denoised-chimera-free SVs were re-assigned using the SILVA 132 taxonomy using qiime2 v2021.11, rescript, and biom. This was done by using rescript to download the SILVA 132 SSU nr99 sequences and taxonomy files, culling sequences using the default settings (5+ ambiguous bases and homopolymers 8+ bp in length), filtering out short sequences using the default settings (removing archaea < 900 bp, bacteria < 1200 bp, and eukaryote < 1400 bp), dereplicating the sequences using p-mode ‘uniq’. A 16Sv4v5 amplicon-region specific classifier was created by extracting this region using the 515f and 926r primers, using rescript to dereplicate the extracted sequences using p-mode ‘lca’ and p-perc-identity 0.97 then evaluating and fitting a naive Bayes classifier. Using the SILVA 132 dataset, this classifier had the following metrics: precision 0.95 (species) - 1 (Domain), recall 0.87 - 1, and F-measure 0.91 - 1. Our denoised-chimera free SVs were imported into qiime and classified. These taxonomic assignments were combined with our SV table (SVs x samples) and used with FAPROTAX to assign function to our 16S SVs. Unassigned SVs were labeled ‘Unknown’. To try to improve the annotation of terrestrial bacteria, additional annotations for nitrogen-fixers, potassium- and phosphate-solubilizers, and chitinolytic bacteria were added.^115–118^

Fungal taxonomic assignments at the class rank were estimated to be at least 95% correct. Genera were retained if bootstrap values were at least 0.70 for assignments that were at least 80% correct. Species were retained if the bootstrap values were at least 0.80 for assignments that were at least 70% correct based on leave one sequence out testing using a reference sequence database based on UNITE and assuming that query taxa are present in the reference set.^103^ ITS SVs were screened for the presence of known pyrophilous (fire-loving) fungi based on studies that used sporocarp sampling at the genus and species ranks.^43,47,76^ Fungal ITS sequences were then assigned using the FungalTraits database by importing their reference set into R.^119^ Functional assignments (trophic_mode_fg) were assigned to our SVs at the genus rank. Unassigned SVs were labeled ‘Unknown’. We attempted to improve the guild & trait annotations from FungalTraits by adding additional information from the literature for rhizomorph-forming, ericoid, and fire-associated fungi (based on sporocarp sampling).

Arthropod taxonomic assignments at the class rank were estimated to be 99% correct. Genera were retained if the bootstrap value was at least 0.30 for assignments that are at least 99% correct. Species were retained if the bootstrap support value was at least 0.70 for assignments that were at least 95% correct based on leave one sequence out testing and assuming that the query taxa are present in the refence set.^31^ Arthropod SVs at the family rank were assigned function from several databases. Fungal feeding groups (6 categories: collector-filterer (CF), collector-gatherer (CG), herbivore (scraper) (HB), shredder (SH), predator (piercer, engulfer), parasite (PA)) were obtained from the United States Environmental Protection Agency (EPA) Freshwater Biological Traits Database available from https://www.epa.gov/risk/freshwater-biological-traits-database-traits.^120^ Feeding types according to Moog, 1995 and feeding habits according to Tachet et al., 2010 were obtained from the freshwaterecology.info database and re-coded to match the EPA functional feeding groups plus ‘Other’ for other feeding types.^121–123^ At the family rank, it is sometimes possible to assign multiple functions, so these were concatenated into their own functional categories. Unassigned SVs were labeled ‘Unknown’. Note that these macroinvertebrate trait databases are largely focused on aquatic species. To improve the proportion of terrestrial species annotated, we broadly annotated the Arachnida to the order rank; the Collembola were broadly annotated at the class rank; and the Hymenoptera were annotated to the family rank.^124–128^ COI species and genera were screened for the presence of known pyrophilous (fire-loving) insects.^51–53^

We used the Pearson’s phi coefficient of association with binary data to assess the strength of association between potential bioindicator taxa and development stage in fire-origin sites using the multipatt function from the ‘indicspecies’ package.^129^ We set the multipatt function to ‘r.g’ to correct for groups that may have more samples than others, with duleg set to TRUE to avoid considering combinations of groups, with 9999 permutations. We only retained potential bioindicators if the p-value was < 0.05.

Nestedness based on overlap and decreasing fill (NODF) was calculated using the nestednodf algorithm with the oecosimu function with 999 replicates to generate a null distribution to assess significance.^61^ Specialization, or the extent that the stand development stage ‘selects for’ sequence variants, was assessed with the network-wide H_2_′ index using the H2fun function from the bipartite package and significance was assessed by comparison with a null distribution based on 999 randomized matrices generated with the r2dtable function.^130^

We simulated the truncation of stand development stages due a decreased fire return interval. We assessed the effect of progressively removing the oldest stand successional stages on gamma diversity to assess the effect on landscape diversity and on NODF to assess the effect on community nestedness and stability.

## Acknowledgements

TMP was funded by the Government of Canada through the Genomics Research and Development Initiative (GRDI) Metagenomics-Based Ecosystem Biomonitoring (Ecobiomics) project. We are grateful to Susan Bowman at the Great Lakes Forestry Centre for soil sample processing and conducting DNA extractions; Marie-Josée Morency and the Seguin lab for amplicon and sequencing library preparation; as well as to the Natural Research Council Canada sequencing facility in Saskatoon for Illumina MiSeq sequencing services.

## Author contributions

DM and LV conceived of and obtained funding for the project. TMP did the bioinformatics, data analysis, and wrote the first draft of the manuscript. ES, DM, and LV assisted with data analysis. All authors contributed to editing the manuscript.

## Data availability

Sequences have been deposited to the NCBI SRA under the GRDI-Ecobiomics project accession PRJNA565010 for the BioSample accessions SAMN26926703 - SAMN26926795 used in this study and data will be released upon manuscript acceptance. The MetaWorks v1.9.3 multi-marker metabarcode bioinformatic pipeline is available from https://github.com/terrimporter/MetaWorks. The COI classifier trained to work with the RDP Classifier is available from https://github.com/terrimporter/CO1Classifier. The ITS classifier based on the ITS UNITE+INSD full dataset v8.2 and trained to work with the RDP Classifier is available from https://github.com/terrimporter/UNITE_ITSClassifier. Infiles and scripts used to generate manuscript figures are available along with other digital resources including: 1. A list of fire-associated arthropods largely from Wikars (1997). 2. A list of fire-loving fungi largely from Zak and Wicklow (1980) and Dix and Webster (1995). 3. A list of terrestrial arthropod functional feeding guilds meant to facilitate comparisons with freshwater macroinvertebrate FFGs. 4. A list of fungal traits for rhizomorph-forming, ericoid, and fire-associated fungi that can be used to supplement FungalTraits annotations. 5. The Sansupa et al., 2021 enhanced FAPROTAX annotations further supplemented with chitinolytic taxa is available here in a digital format; available on GitHub from https://github.com/terrimporter/Chronosequence.

## Notes

### Competing Interest Statement

The authors have declared no competing interest.

### Summary of Updates

Clarified methods. New Fig 1.

