## Supplementary Material for "All boreal forest successional stages needed to maintain the full suite of soil biodiversity, community composition, and function following wildfire"

**Supplementary Results**

**Fig S1. Fire-origin soil communities are largely distinct from one stand development stage to the next.** Pairwise permutational analysis of variance (PERMANOVA) Holm adjusted p-values are shown for each successive pairs of stand development stages. Results are shown for analyses conducted at the a) sequence variants level (homogeneous variances), b) genus (homogeneous variances, except for arthropoda), and c) family ranks (homogenous variances, except for bacteria). Abbreviations: bacteria (B), fungi (F), arthropoda (A); establishment (E), crown closure (CC), thinning (T), late self-thinning (LST), and mature (M) stand development stages.

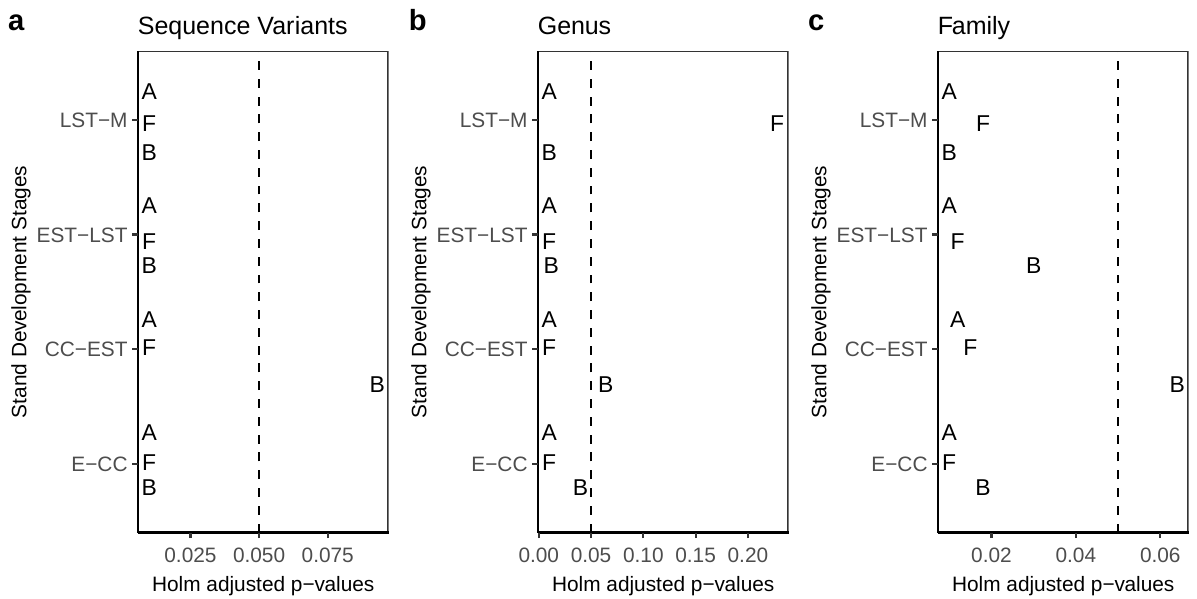

Because we were able to annotate taxonomy and function to finer taxonomic ranks for fungi, we looked more closely at the nestedness and specialization patterns for the dominant fungal taxonomic and functional groups (Fig S2). Some functional groups such as undefined saprotrophs and ectomycorrhizal fungi, as well as all the major fungal taxonomic groups were significantly more nested than null expectations (NODF). The nestedness pattern in ectomycorrhizal fungi in particular, show that relatively few sequence variants are shared across all stand development stages, ectomycorrhizal fungi in this study have a relatively small core community, and a large number of SVs come and go between stages in a nested pattern consistent with what we see for fungi overall (described above) (Fig S3).

**Fig S2. Fungal nestedness and specialization across stand development stages.** We compare community composition of the most abundant taxonomic and functional groups based on: a) nestedness based on overlap and decreasing fill (NODF), and we assess the extent that stand development stages ‘select for’ species using the b) network-wide H_2_′ index. NODF is based on the presence-absence data and the H_2_′ index is based on the relative read abundance of sequence variants. Groups that are more nested (NODF) or specialized (H_2_′ index) than null expectations are indicated with asterisks: p-value < 0.05 (*), < 0.01 (**), or < 0.001 (***).

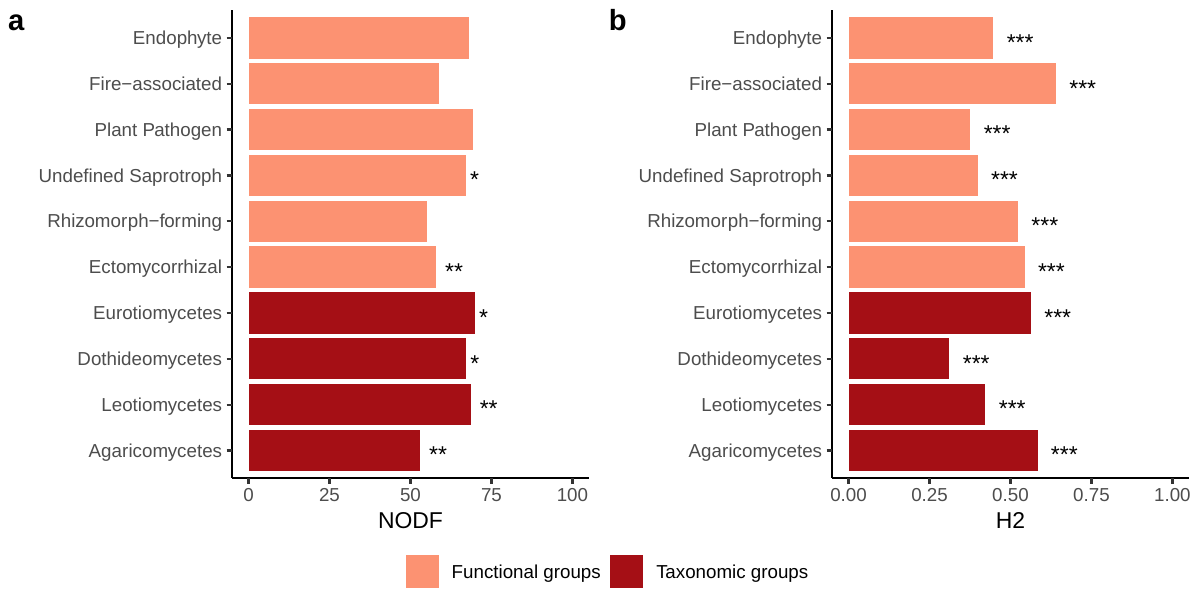

The extent that stand development stages ‘select for’ taxa was measured using the network-wide H_2_′ index. The H_2_′ index is scaled from 0 (generalized) to 1 (specialized). Network-wide specialization was significantly higher than null expectations, but still relatively low overall (bacteria 0.07; fungi 0.20; arthropoda 0.19, p-value < 0.001 for each), especially for bacteria where many of the same sequence variants were detected at each development stage. Network-wide specialization (H_2_′) was higher than null expectations for all major fungal functional and taxonomic groups, but especially for fire-associated fungi (64%)(Fig S2b). These results are supported by the large number of stand development stage bioindicators we detected.

**Fig S3. Nestedness patterns vary across major fungal functional and taxonomic groups.** Nestedness based on overlap and decreasing fill (NODF) are shown below. Nestedness is measured on a scale from 0 (random) to 100 (perfectly nested). Grey bars indicate the presence of a sequence variant (SV), and a white bars indicates the absence of a SV. Grey bars stacked from the left represent the proportion of the community shared across development stages, the core community, and towards the right, reflects changing SV occupancy across development stages. Grey bars stacked from the top to bottom represents nestedness from the stand development stages with the greatest richness to stages with the lowest richness.

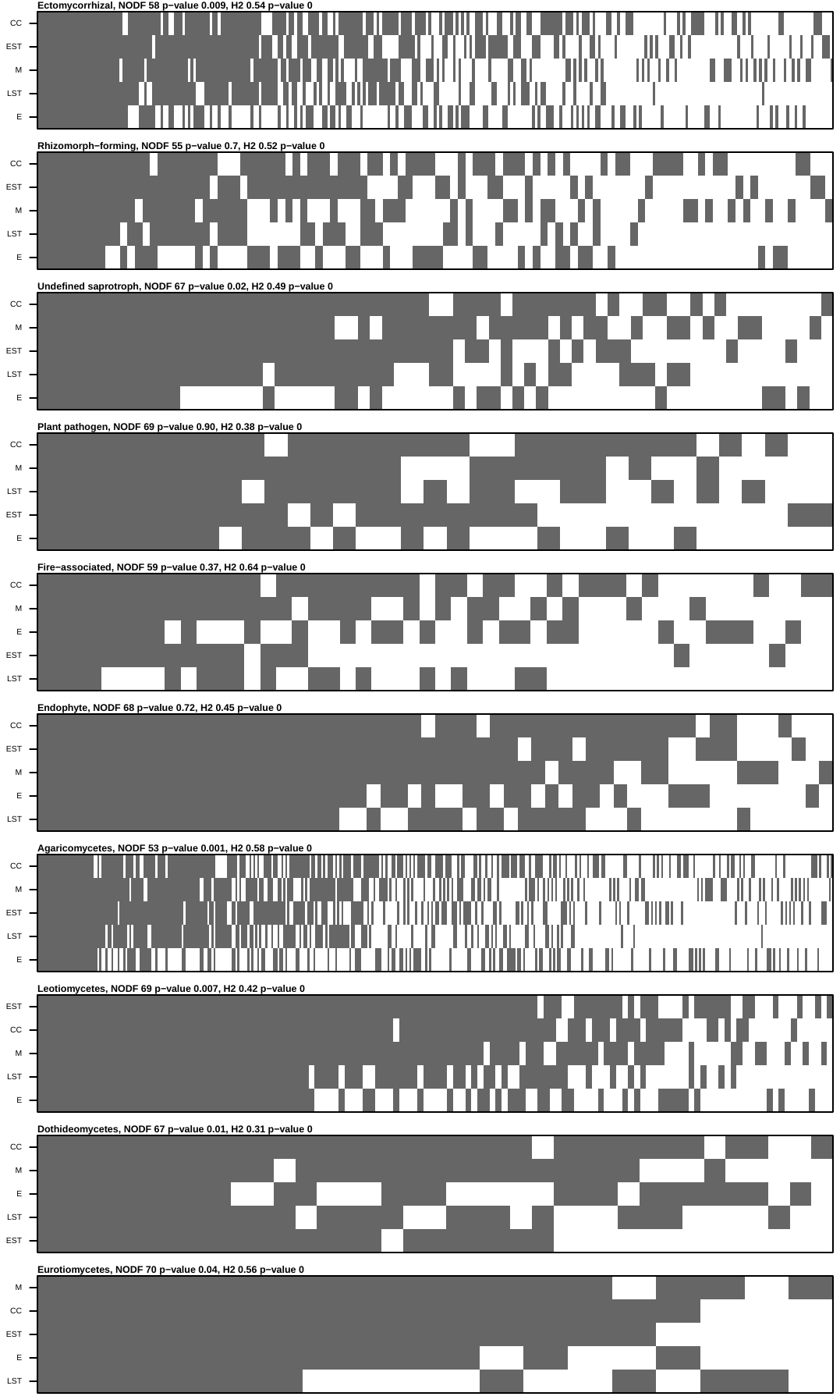

**Fig S4. The diversity and frequency of fire-associated fungal species is highest from fire-origin sites at the establishment stage.** Results are shown at the a) species hypothesis and b) genus levels. Frequency is the number of unique sequence variants per sample per development stage. Abbreviations: establishment (E), crown closure (CC), early self-thinning (EST), late self-thinning (LST), mature (M).

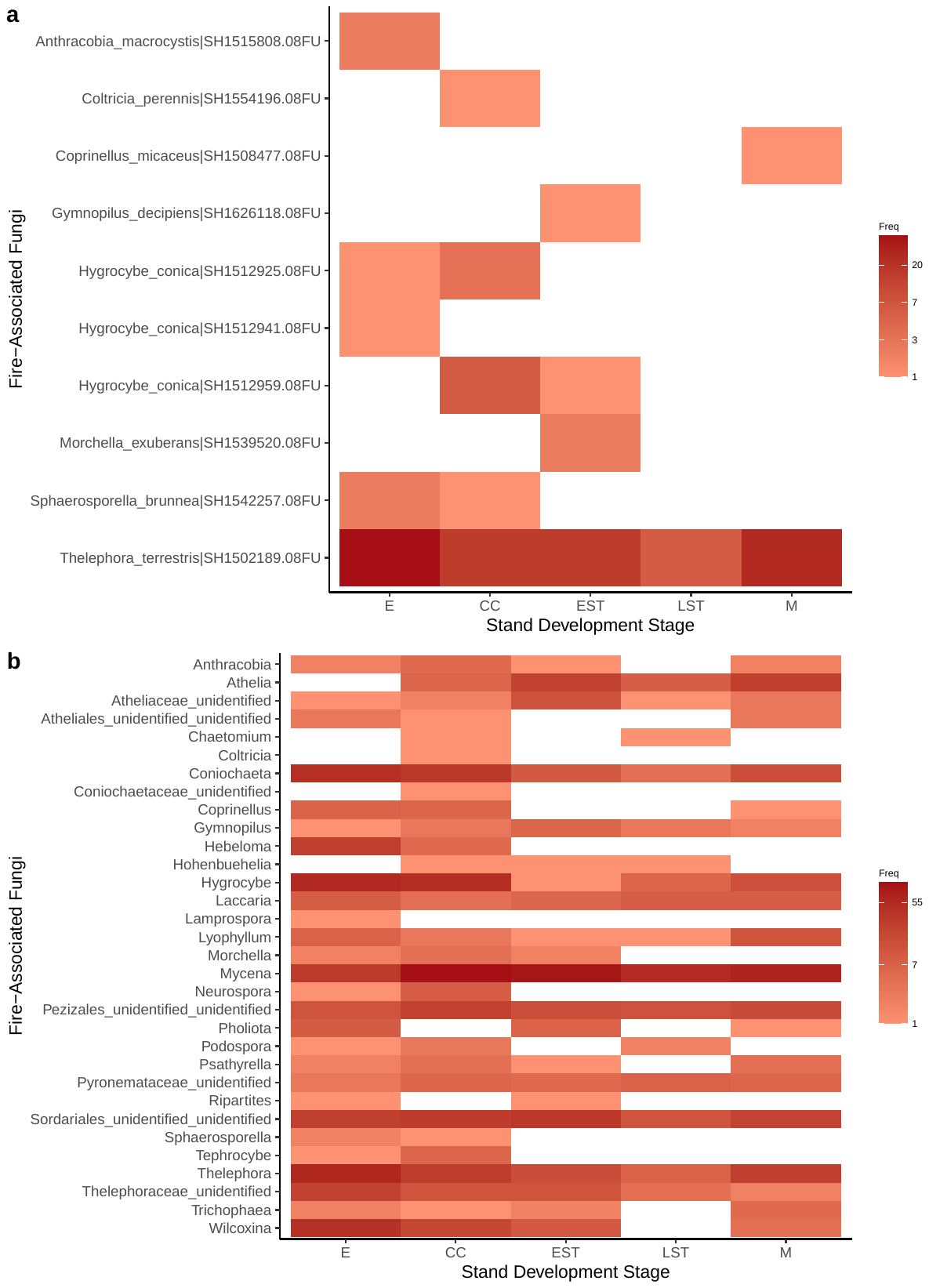

For bacteria, we checked for known fire-associated bacteria at the genus rank as this was the finest level of resolution we could get with confidence using the 16S RDP classifier (Fig S5). Fifteen out of 134 (11%) fire-associated bacterial genera known from previous studies were not represented in our reference sequence database. At this taxonomic resolution, many genera appear to be cosmopolitan, such as *Aquicella*, *Ferruginibacter*, and *Sphingomonas* as they are found across many sites. It is also likely that these genera contain species that may not be preferentially associated with post-fire habitats and a finer level of taxonomic resolution would be preferred when assigning bacterial ecological function.

**Fig S5. Frequency of bacterial genera that may be associated with fire-origin sites.** Frequency (Freq) is the sum of unique sequence variants per sample per development stage. Abbreviations: establishment (E), crown closure (CC), early self-thinning (EST), late self-thinning (LST), mature (M).

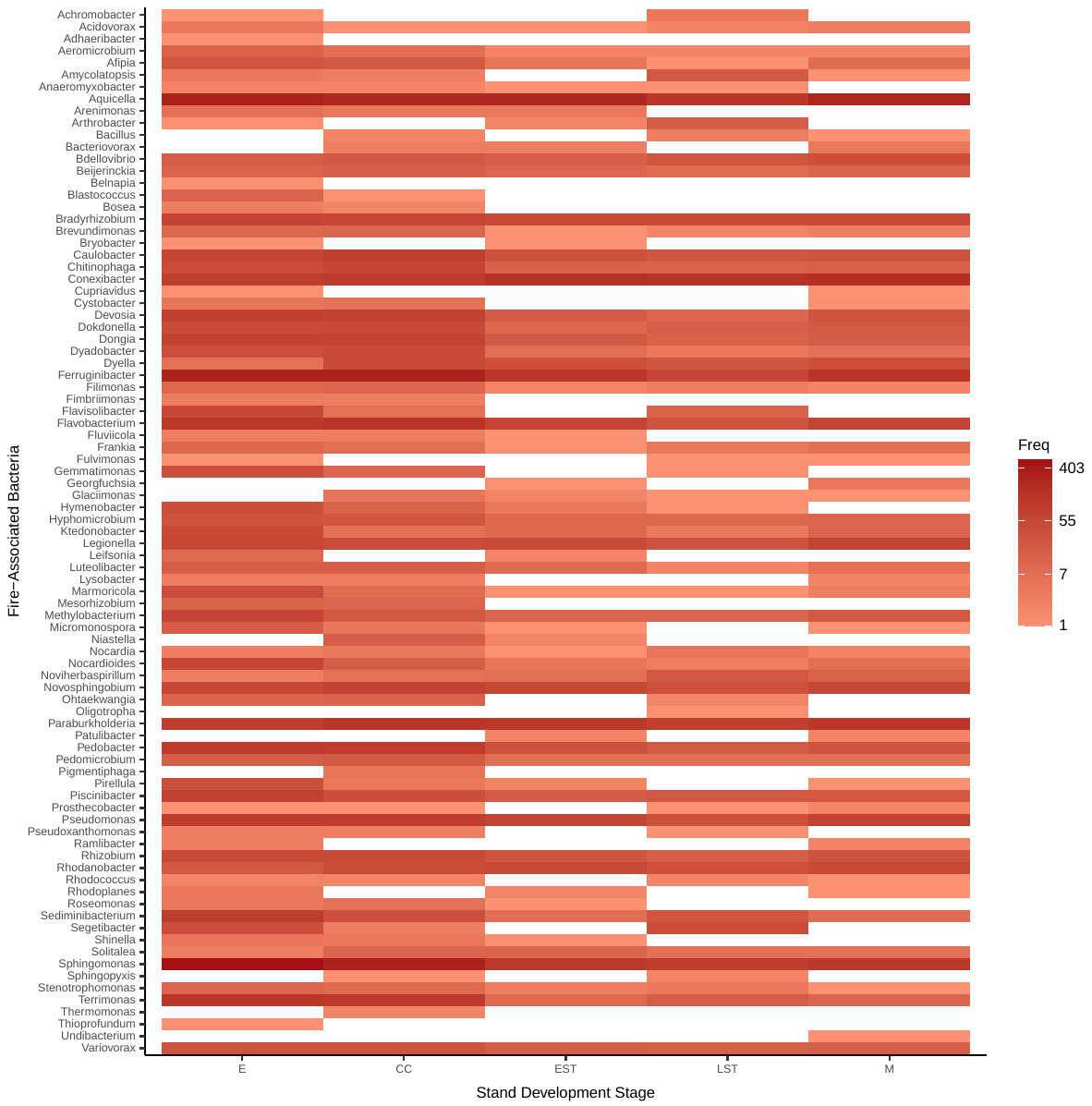

For arthropods, we did not detect any fire-associated insect species known from previous studies. Because 39 out of 66 (59%) previously known fire-associated insect species were not represented in our reference sequence database, we instead summarized our results at the genus rank where only 7 out of 41 (10%) targeted fire-associated insect genera were missing (Fig S6). At this taxonomic resolution, many genera such as *Agonum*, *Pterostichus*, *Quedius*, and *Stegana* appear cosmopolitan across each stand development stage. It is also likely that these genera contain species that are not strictly associated with post-fire habitats and a species level resolution would be preferred for assigning ecological function along with a reference sequence database that better represents fire-associated arthropods.

**Fig S6. Frequency of insect genera that may be associated with fire-origin sites.** We did not detect any species of fire-loving insects. Here we present the frequency of genera associated with known fire-loving insect species. Frequency is the number of unique sequence variants per sample per development stage.

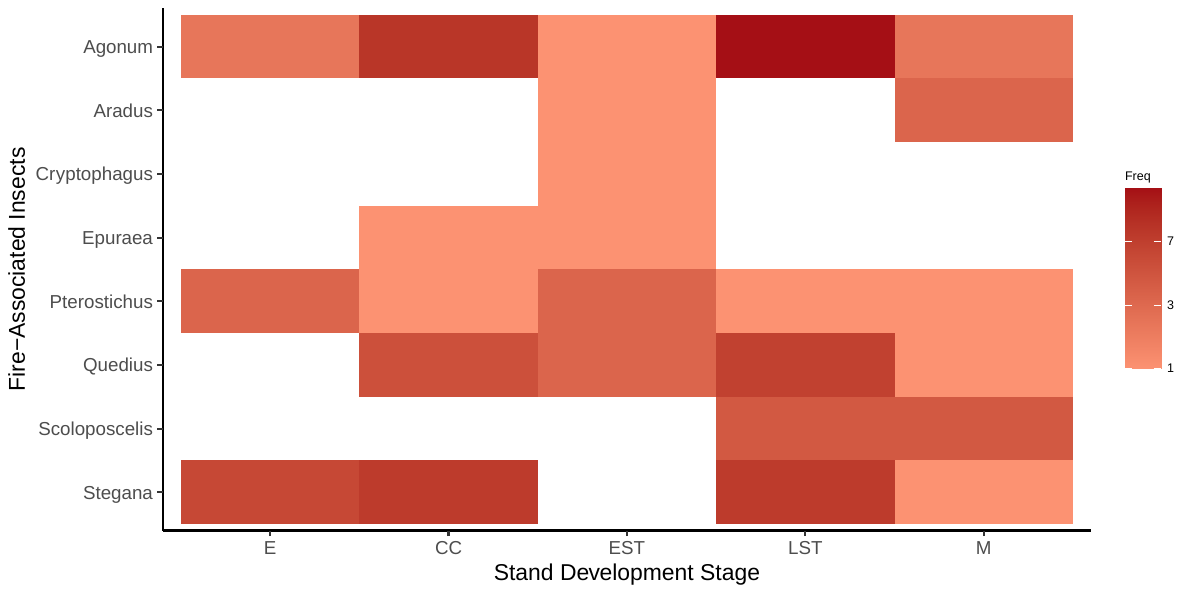

**Fig S7. Rarefaction curves plateau indicating sequencing depth was sufficient for each amplicon.** Across all samples in this study, the number of sequence variants (SVs) detected from bacteria (16S) is highest, followed by SVs from fungi (ITS), and least for SVs from arthropods (COI).

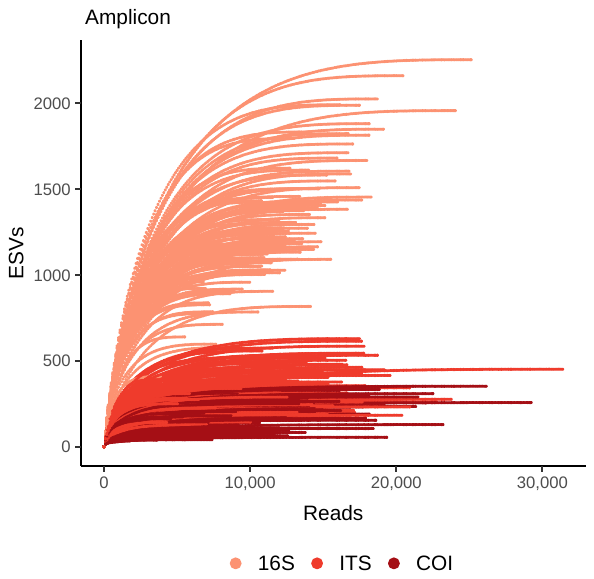

**Table S1.** **Fire-associated fungal taxa detected in this study: 8 species (10 SHs) and 24 additional taxa that may also represent fire-associated fungi.**

| **Genus** | **Specific Epithet** | **Species Hypothesis** |
| --- | --- | --- |
| *Anthracobia* | *macrocystis* | SH1515808.08FU |
| *Coltricia* | *perennis* | SH1554196.08FU |
| *Coprinellus* | *micaceus* | SH1508477.08FU |
| *Gymnopilus* | *decipiens* | SH1626118.08FU |
| *Hygrocybe* | *conica* | SH1512925.08FU |
| *Hygrocybe* | *conica* | SH1512959.08FU |
| *Hygrocybe* | *conica* | SH1512941.08FU |
| *Morchella* | *exuberans* | SH1539520.08FU |
| *Sisporaphaerosporella* | *brunnea* | SH1542257.08FU |
| *Thelephora* | *terrestris* | SH1502189.08FU |
| *Athelia* |  |  |
| Atheliaceae unidentified |  |  |
| Atheliales unidentified |  |  |
| *Chaetomium* |  |  |
| *Coniochaeta* |  |  |
| Coniochaetaceae unidentified |  |  |
| *Hebeloma* |  |  |
| *Hohenbuehelia* |  |  |
| *Laccaria* |  |  |
| *Lamprospora* |  |  |
| *Lyophyllum* |  |  |
| *Mycena* |  |  |
| *Neurospora* |  |  |
| Pezizales unidentified |  |  |
| *Pholiota* |  |  |
| *Podospora* |  |  |
| *Psathyrella* |  |  |
| Pyronemataceae unidentified |  |  |
| *Ripartites* |  |  |
| Sordariales unidentified |  |  |
| *Tephrocybe* |  |  |
| Thelephoraceae unidentified |  |  |
| *Trichophaea* |  |  |
| *Wilcoxina* |  |  |

**Table S2. [Excel spreadsheet] Stand development stage bioindicators from fire-origin sites.** Bioindicator taxa are shown for bacteria, fungi, and arthropods if the Pearson’s phi coefficient of association was significant (p-value < 0.05). Abbreviations: p-value < 0.05 (*), < 0.01 (**), < 0.001 (***).
